# Artificial microRNAs targeting Tau enable post-symptomatic functional recovery in aged tauopathy mice

**DOI:** 10.1101/2025.10.14.680333

**Authors:** Carolina Lucía Facal, Indiana Páez-Paz, A. Ezequiel Pereyra, Ramiro Clerici-Delville, Clara Gaguine, Rocío Foltran, Mariano Soiza-Reilly, María Elena Avale

## Abstract

Tauopathies are a group of neurodegenerative disorders, including Alzheimer’s disease, frontotemporal dementia, and progressive supranuclear palsy, characterized by the pathological accumulation of tau protein. While tau reduction has emerged as a promising disease-modifying strategy, most preclinical studies have focused on preventive approaches, and the therapeutic potential after clinical onset remains largely unexplored. This limitation is critical, as patients are typically diagnosed after symptoms emerge. Furthermore, global tau suppression may disrupt physiological tau functions and lead to adverse effects, underscoring the need for targeted interventions.

RNA-based therapies, particularly microRNA (miRNA)-mediated silencing, offer high specificity, versatility, and sustained target knockdown. Here, we developed artificial microRNAs (Tau-miRNAs) designed for site-directed expression to selectively reduce tau levels in vulnerable brain regions, thereby minimizing off-target effects. We tested the efficacy of Tau-miRNAs in a tauopathy mouse model at advanced disease stages, delivering them into the prefrontal cortex after cognitive and electrophysiological deficits had developed. This post-symptomatic intervention led to long-term improvements in memory, restoration of neuronal firing properties, and reduced pathological tau at synapses. Our findings highlight the therapeutic potential of spatially targeted RNA-based tau-lowering strategies for late-stage intervention in tauopathies, addressing a critical unmet need in the treatment of these devastating disorders.

## INTRODUCTION

Tauopathies are a group of neurodegenerative diseases characterized by the pathological accumulation of tau protein in certain brain nuclei, leading to synaptic deficits, abnormal neuronal activity, and ultimately neurodegeneration^1,2^. These pathologies include Alzheimer’s disease (AD), progressive Supranuclear Palsy (PSP) and frontotemporal dementia (FTD), which have devastating cognitive and behavioral effects.

In the healthy brain, Tau plays a myriad of physiological functions, maintaining microtubule dynamics and facilitating intracellular transport, both of which are essential for neuronal function^3,4^. However, several failures in genetic or metabolic pathways are known to alter tau function leading to the accumulation of pathological tau variants, including hyperphosphorylated, misfolded, or truncated forms^5,6^. Under these pathological conditions tau dissociates from the axon and relocates to the somatodendritic compartment where it exerts detrimental changes in neuronal function ^7–9^, assembling into insoluble filaments and finally forming neurofibrillary tangles (NFTs) ^6,7,10^. Such mislocalization underlie severe neuronal impairments that could begin years before neuronal death^10,11^. Although NFTs correlate with the progression of the disease in a discrete pattern that goes from the primary affected areas to the spreading of pathological tau that leads to synapse loss and neurodegeneration^12^, evidence suggests that the “gain of toxic function” of tau is less attributed to the highly assembled structures like NFTs, and more related to the previous oligomeric forms^13,14^, which are abundantly found in synaptic terminals of patients ^7^.

In this context, disease-modifying therapies that lower the levels of intracellular pathological tau species upstream of protein aggregation may prove effective in reversing the pathological mechanisms associated with tauopathies^5,15,16^. Noteworthy, reducing or ablating tau can confer protective effects in preclinical models of neurodegeneration, including mitigating excitotoxicity and enhancing synaptic function ^17–20^. Yet, most preclinical studies that validate such interventions so far have focused on preventive strategies, limiting the scope of the therapeutic window. Developing more realistic and efficient therapies would require targeting tau after the onset of clinical symptoms underlying disease.

As a proof of concept to validate tau synthesis inhibition as a therapeutic strategy, we evaluated the effect of localized tau reduction on reversing cognitive impairments in a mouse model of tauopathy after symptom onset. By intervening post-onset, we aimed to assess the feasibility of phenotype rescue in a context that better mimics clinical scenarios. To achieve this, we utilized a molecular tool that suppresses tau expression through RNA interference. Specifically, we developed artificial tau-targeting microRNAs (Tau-miRNAs), which have previously demonstrated efficient target engagement and tau reduction in both human neurons and mouse brain ^21^. These Tau-miRNAs were expressed using lentiviral vectors, ensuring stable, long-term expression after a single delivery. We performed administration of tau-miRNAs into the prefrontal cortex of aging htau mice after detecting cognitive decline and impairments in neuronal activity. Cognitive deficit was assessed in novel object recognition tasks, followed by field records to determine neuronal firing patterns in prefrontal neurons. Postmortem analyses showed reduction of pathological Tau accumulation, while array tomography showed a marked decrease of phospho-Tau clusters at presynaptic terminals after Tau-miRNA treatment.

Our findings provide insights into the therapeutic potential of tau-miRNA and its ability to reduce pathological tau accumulation, rescue neuronal activity and cognitive function after phenotypic onset.

## RESULTS

### Local tau reduction in the prefrontal cortex rescues cognitive impairments and modulates neuronal firing rate in aged htau mice

Based on our previous observation that local injection of Tau-miRNA effectively prevents pathological tau accumulation in htau mice ^21^, we wondered whether tau reduction shortly after phenotypic onset could rescue or reverse pathological phenotypes in this model. Htau mice accumulate hyperphosphorylated and insoluble tau predominantly in the prefrontal cortex during aging^22,23^. In fact, FDG-PET analyses have shown progressive metabolic decline in the prelimbic area of the prefrontal cortex (PFC) between 3 and 12 months of age^24^. From 6 months onward, htau mice exhibit cognitive decline as evidenced by deficits in the novel object recognition task ^24^. Additionally, in vivo electrophysiological field recordings reveal exacerbated neuronal firing rates in the PFC of aging htau mice^21^. To evaluate the therapeutic potential of tau reduction in symptomatic mice, we performed a single administration of Tau-miRNAs into the PFC of htau and wild-type (WT) mice at 6 months of age, when deficits associated with prefrontal dysfunction are already established.

Six-month-old wild-type (WT) and htau mice underwent baseline behavioral testing prior to injection to assess cognitive decline in htau mice relative to WT controls and then were randomly assigned to treatment groups to receive a single injection of either Tau-miRNAs or scrambled control (Scr) miRNAs into the medial prefrontal cortex (mPFC). Behavioral, electrophysiological, and molecular analyses were performed again at 12 months of age (6 months post-injection) to evaluate phenotypic rescue (Figure 1A). At 12 months, htau mice treated with scrambled miRNA (Scr) showed significant cognitive impairment in the novel object recognition (NOR) task, with an average discrimination index (DI) of approximately 50%, indicating impairment to distinguish the novel from the familiar object. In contrast, Tau-miRNA-injected htau mice demonstrated a robust preference for the novel object, with a mean DI around 70%, comparable to WT littermates (Figures 1B-C and S1A). Importantly, total exploration time did not differ between groups (Figures S1A-B), and no sex differences in DI were observed (Figure S1C), suggesting both males and females respond similarly to Tau-miRNA treatment. Within-group longitudinal analysis revealed that all htau mice receiving Tau-miRNA improved their NOR performance post-injection, whereas most htau mice injected with Scr-miRNA exhibited worsened cognitive performance over time (Figure 1D).

**Figure 1.**
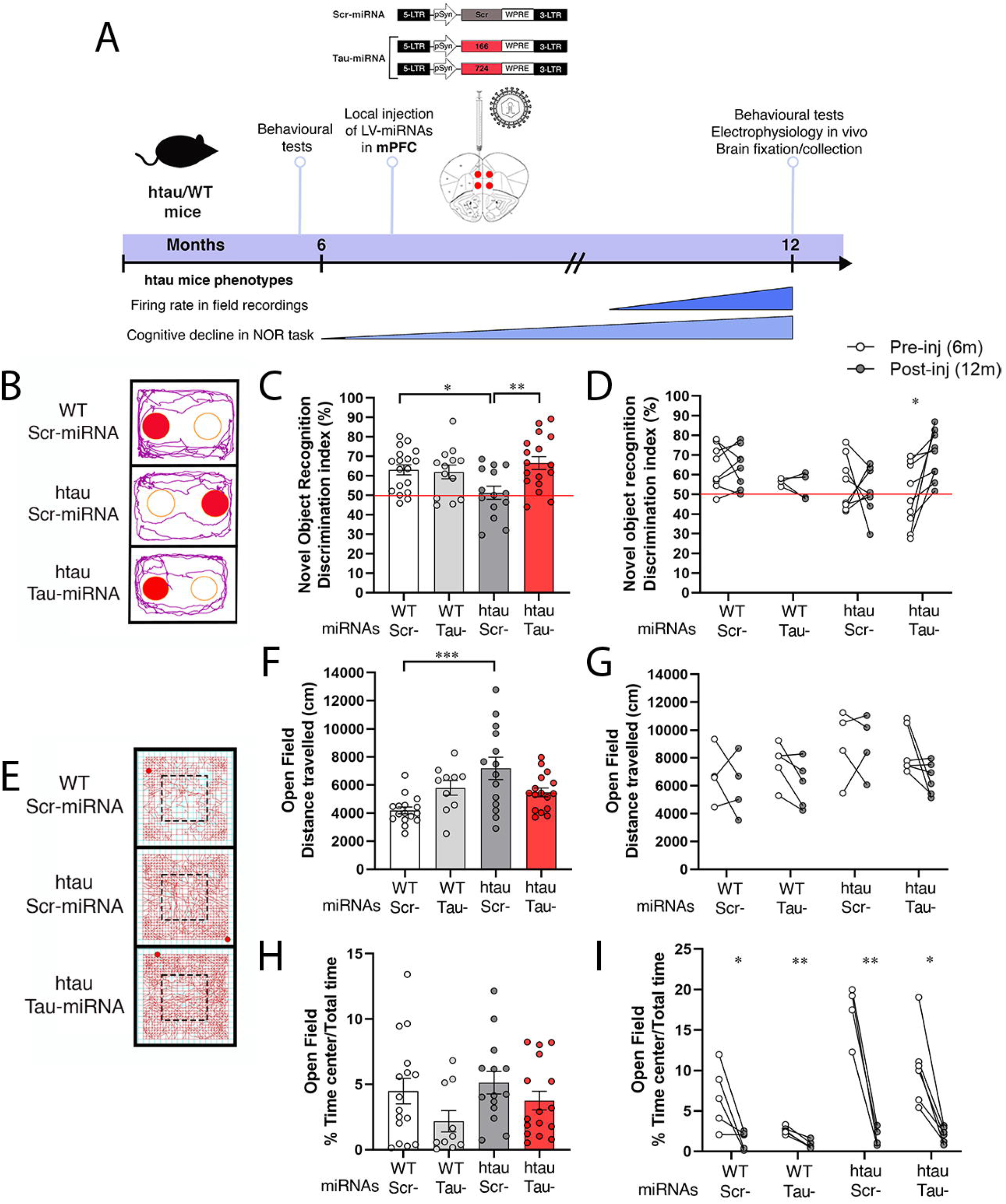
Single injection of Tau-miRNA in the PFC specifically rescues cognitive decline in aged htau mice. **A**. Timeline of lentiviral injection of Tau-miRNA in the mPFC of htau mice. 6-months-old mice (htau and WT littermates) were injected after phenotypic onset related to pathological tau accumulation in the PFC of htau mice. LVs contained either a scrambled control miRNA or an equimolar combination of two artificial microRNAs targeting *MAPT* mRNA. After injection, mice were maintained until 12 months of age, where behavioural, electrophysiological and molecular analyses were performed. Time course of some pathological phenotypes related to tau accumulation in htau mice are described such as beginning of cognitive decline in the novel object recognition (NOR) task at 6 months old and the increasement of firing rates of neurons in the PFC described by electrophysiological field recordings *in vivo*. **B-D**. NOR performance in miRNA-injected groups. **B**. Representative traces of exploration, where circles describe the area for the novel (filled with red) and familiar (empty) objects. Discrimination index in the NOR task for each group (**C**) analysed at 12 months of age and (**D**) comparing each animal before (Pre-inj 6m) and after (Post-inj 12m) the injection. For Figure C: WT Scr-miRNA=18, WT Tau-miRNA=13, htau Scr-miRNA=14 and htau Tau-miRNA=17; and D: WT Scr-miRNA=8, WT Tau-miRNA=4, htau Scr-miRNA=7, htau Tau-miRNA=9. **E-I**. Open field task in miRNAs-injected groups. **E**. Representative traces of exploration (in red). Total distance travelled for each group (**F**) assessed at 12 months and (**G**) before and after treatment. Time spent in the center relative to total time of exploration for each group (**H**) analysed at 12 months and (**I**) before and after the injection. For Figures F and H: WT Scr-miRNA=16, WT Tau-miRNA=10, htau Scr-miRNA=14, htau Tau-miRNA=16; and G and I: WT Scr-miRNA=4/5, WT Tau-miRNA=5, htau Scr-miRNA=4, htau Tau-miRNA=6. For Figures C, F and H: data is shown as scatter dot plots with bars ± SEM, One-way ANOVA followed by Tukey’s *post hoc* test; and D, G and I: data is shown as scatter dot plots, paired *t*-test. *p*\*<0,05, *p*\*\*<0,01 and *p*\*\*\*<0,001.

To assess hyperactivity, another phenotype previously described in the htau model ^22,24,25^, mice were also tested in the open field arena (Figures 1E–G). At 12 months of age, htau mice injected with scrambled miRNA (Scr) displayed significantly increased spontaneous locomotor activity compared to WT controls (Figure 1F), consistent with prevoius analyses^22,24,25^. In contrast, Tau-miRNA-injected htau mice exhibited locomotor activity levels comparable to those of WT mice (Figures 1F–G), indicating normalization of this phenotype upon tau reduction in the PFC. Notably, no differences were observed between groups in the time spent in the center of the open field arena (Figures 1H–I) or in the time spent in the open arms of the elevated plus maze (Figures S1D–E) at 12 months of age. However, both parameters were elevated in htau mice at 6 months of age compared to WT littermates (Figure 1I and Figure S1F), consistent with our previous findings indicating that behavioral disinhibition is a prominent trait of young htau mice that diminishes with age^24^.

We next assessed whether Tau-miRNA treatment could also normalize dysregulated neuronal activity in htau mice. In vivo extracellular field recordings were performed in the medial prefrontal cortex (mPFC) of 12-month-old mice. Recorded neurons were classified as putative pyramidal cells or interneurons based on their mean spike waveform (Figures 2A and S2A), and their firing and bursting activity were analyzed. In line with previous findings showing increased cortical pyramidal firing rates in aging htau mice correlating with cognitive impairment, control htau mice injected with Scr-miRNA exhibited elevated firing rates (Figures 2B–C) and increased burstiness (Figure 2D) in pyramidal neurons compared to WT mice. In contrast, Tau-miRNA-injected htau mice displayed significantly lower pyramidal firing rates and burst activity, reaching levels comparable to age-matched WT controls (Figures 2B–D).

**Figure 2.**
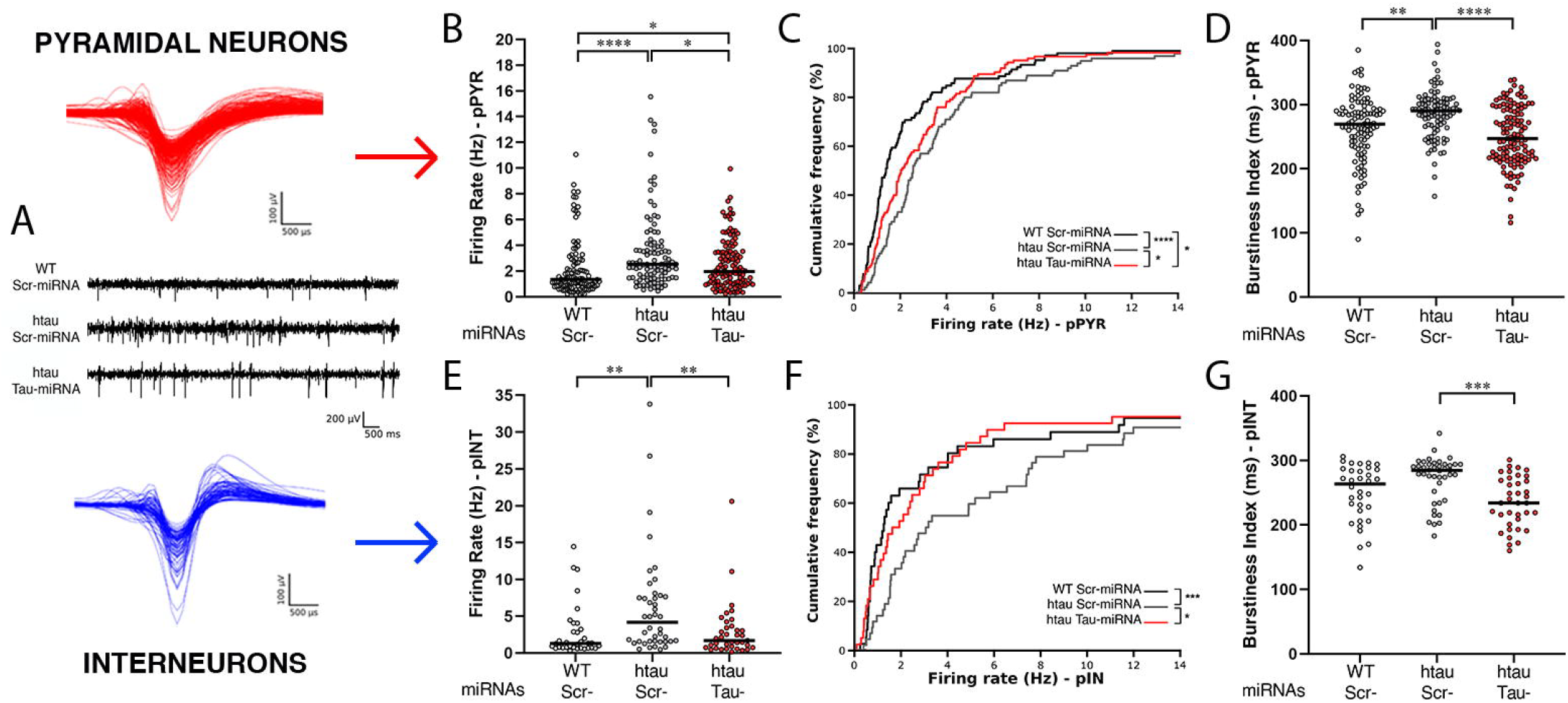
Tau-miRNA directed expression modulates dysregulated firing and bursting activity of cortical neurons in the PFC of htau mice. **A**. *Middle:* Representative traces of an electrophysiological signal band pass-filtered between 300 and 6,000Hz for miRNA-injected groups. Putative pyramidal neurons (*up*, red) or putative interneurons (*bottom*, blue) were classified upon their average spike waveforms. Mice: WT Scr-miRNA=8, htau Scr-miRNA=8 and htau Tau-miRNA=10. **B-D**. Firing rate (**B**), cumulative frequencies (**C**) and burstiness index (**D**) of putative pyramidal neurons (pPYR) registered in the mPFC of injected groups. WT Scr-miRNA=106, htau Scr-miRNA=100 and htau Tau-miRNA=123. **E-G**. Firing rate (**E**), cumulative frequencies (**F**) and burstiness index (**G**) of putative interneurons (pINT) registered in the mPFC of injected groups. WT Scr-miRNA=34, htau Scr-miRNA=42 and htau Tau-miRNA=37. Each dot represents the mean firing rate (Figures B and E) or the burstiness index (Figures D and G) of each recorded neuron along the session, black lines indicate the median value per group and statistical analyses were performed by Kruskall-Wallis followed by Dunn’s *post hoc* test. For Figures C and F, statistical analyses were performed by Kolmogorov-Smirnov test. *p*\*<0,05, *p*\*\*<0,01, *p*\*\*\*<0,001 and *p*\*\*\*\*<0,0001.

Interestingly, a similar pattern was observed in interneurons. Scr-injected htau mice showed elevated interneuron firing rates compared to WT controls (Figures 2E–F), while Tau-miRNA treatment reduced interneuron excitability to WT-like levels (Figures 2E–G). Likewise, both pyramidal neurons and interneurons from htau Scr-miRNA controls exhibited an increased bursting index (Figures S2G–K), suggesting greater propensity for bursting events, which was normalized by Tau-miRNA treatment. Furthermore, the distribution of interspike intervals (ISIs) in representative prefrontal neurons revealed distinct firing clusters in Scr-injected htau mice compared to WT and Tau-miRNA-injected htau mice (Figures S2B–D), consistent with the higher burst activity observed in the control group. Together, these findings demonstrate that post-onset tau silencing via Tau-miRNAs effectively normalizes dysregulated firing patterns in both pyramidal neurons and interneurons in the prefrontal cortex of htau mice.

### Tau-miRNA injection reduces insoluble tau and reverts pathological accumulation in synaptic terminals in the mPFC of aged htau mice

We next aimed to determine whether the phenotypic recovery observed after Tau-miRNA treatment correlates with reduction in pathological tau accumulation. Western blot analyses performed with 12-month-old mouse brain extracts revealed that Tau-miRNA administration reduced total tau protein levels by approximately 30% in the injected medial prefrontal cortex (mPFC) of htau mice (Figures 3A–B and S3A), consistent with our previous findings observed after Tau-miRNA administration in presymptomatic - 3 months-old-mice^21^. Moreover, hyperphosphorylated tau, detected with the PHF1 antibody (Figures 3C–D and S3B), and insoluble tau species, assessed by sarkosyl-insolubility assay (Figures 3E–F and S3C), were both significantly decreased in Tau-miRNA-injected htau mice compared to Scr-miRNA controls. These findings indicate that Tau-miRNA delivery after phenotypic onset not only reduces total tau levels, but also diminishes tau pathological burden in its aggregated and phosphorylated forms.

**Figure 3.**
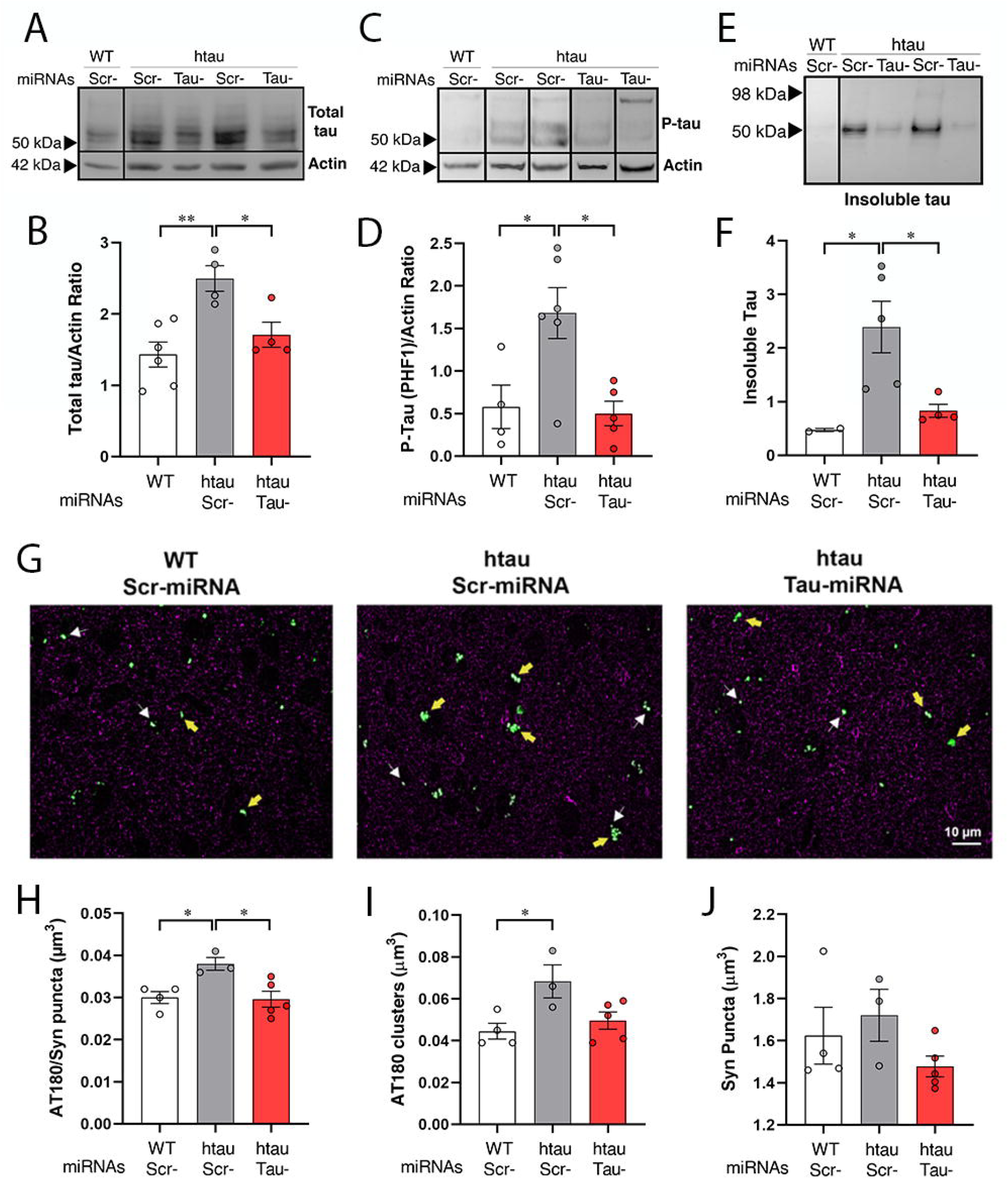
Tau-miRNA reduces insoluble and p-tau levels in the mPFC of htau mice. A-B. **A**) Representative immunoblots and (**B**) quantification of total tau protein contents in the mPFC of miRNA-injected mice. WT=6, htau Scr-miRNA=4 and htau Tau-miRNA=4. WT group represents pooled samples from Scr- and Tau-miRNA injected groups, with similar values obtained for total tau levels. **C-D. C**) Representative immunoblots and (**D**) quantification of p-tau (PHF1) contents in the mPFC of miRNA-injected mice. WT=4, htau Scr-miRNA=6 and htau Tau-miRNA=5. **E-F. E**) Representative immunoblots and (**F**) quantification of insoluble tau contents in the mPFC of miRNA-injected mice. WT Scr-miRNA=2, htau Scr-miRNA=5 and htau Tau-miRNA=4. **G**. Representative images obtained by array tomography, showing p-tau (AT180, green) in the mPFC synaptic neuropil of miRNA-injected mice. Arrows indicate colocalization of AT180 with the presynaptic marker synapsin 1a (magenta). **H-J**. Quantitative analyses of (**H**) density of AT180/synapsin colocalized puncta, (**I**) density of AT180 clusters and (**J**) density of synaptic boutons in the mPFC. WT Scr-miRNA=4, htau Scr-miRNA=3 and htau Tau-miRNA=5. Data is shown as scatter dot plots with bars ± SEM and statistical analyses were performed by One-way ANOVA followed by Tukey’s *post hoc* test. \**p*<0,05 and \*\**p*<0,01.

To further investigate the subcellular distribution of hyperphosphorylated tau in htau mice, we performed high-resolution array tomography immunofluorescence to assess the colocalization of phospho-tau (Thr231) with presynaptic terminals in the mPFC (Figure 3G). As previously described, Scr-injected htau mice exhibited a higher number of p-tau–positive presynaptic terminals compared to age-matched WT controls, along with an increased number of p-tau clusters in the prefrontal cortex (Figures 3G–I). Remarkably, six months after Tau-miRNA administration, p-tau accumulation at synaptic terminals was significantly reduced in htau mice (Figures 3G, right panel; 3H–I, red bars). These results indicate that pathological tau accumulation at the synapse can be effectively diminished following sustained suppression of tau synthesis via Tau-miRNAs. Importantly, the number of synaptic boutons remained unchanged across all groups (Figure 3J), suggesting that the reduction in synaptic p-tau is not secondary to synapse loss.

Taken together, these biochemical and histological analyses demonstrate that a single local administration of Tau-miRNAs in the PFC after phenotypic onset achieves efficient target engagement, leading to a significant reduction in pathological p-tau accumulation.

## DISCUSSION

This study shows that local tau reduction using Tau-miRNA can effectively rescue cognitive impairments, modulate neuronal activity, and reduce pathological tau accumulation in htau mice after phenotypic onset, providing a strong foundation for the development of targeted tau-lowering therapies for post-onset interventions in the treatment of tauopathies. Artificial Tau-miRNAs administered into the PFC of aging htau mice reduced total tau protein levels by approximately 30% in the injected area, consistent with previous study^21^. A significant decrease in insoluble tau species and hyperphosphorylated tau (p-tau) clusters following Tau-miRNA administration further supports its efficacy in mitigating tau pathology even after phenotypic onset. These effects were accompanied by a rescue of cognitive deficits and hyperactivity, aligning with prior studies showing protective effects of tau downregulation in models of excitotoxicity and behavioral dysfunction ^19,26–28^. However, unlike preventive approaches, our study demonstrates the capacity of tau reduction to reverse already-established phenotypes.

Electrophysiological recordings further support these findings, showing that Tau-miRNA treatment normalized the increased firing rate and bursting activity in pyramidal neurons and interneurons in the medial PFC of htau mice. This modulation of hyperexcitability is consistent with the proposed role of tau in regulating neuronal excitability ^29,30^ and expands on previous observations by demonstrating that post-onset tau knockdown can mitigate dysregulated cortical activity^31^. Importantly, array tomography revealed that Tau-miRNA significantly reduced p-tau accumulation at presynaptic terminals—synaptic compartments known to be affected early in tauopathies. The reduction in synaptic p-tau, without changes in synaptic bouton density, suggests that tau-targeting strategies can restore synaptic integrity without compromising synaptic structure.

Together, these findings provide mechanistic support for targeting tau to alleviate tauopathy-related neuronal and synaptic dysfunction. At the molecular level, the observed reduction of insoluble tau and presynaptic p-tau clusters highlights the relevance of tau accumulation at synaptic sites in driving both cognitive and electrophysiological impairments. This aligns with prior work linking tau pathology to synaptic dysfunction and abnormal circuit activity in AD ^7^.

This study underscores the therapeutic potential of targeted, localized tau-lowering strategies to reverse established phenotypes. Since clinical diagnosis often coincides with advanced tau pathology, the ability to rescue function after symptom onset is critical for translational applications. Moreover, the absence of effects on anxiety or general locomotion in treated animals highlights the specificity of local Tau-miRNA intervention and supports its potential safety and tolerability.

Tau reduction targeting *MAPT* mRNA has emerged as one of the most promising therapeutic strategies under clinical evaluation. Several studies have shown that moderate reductions of up to 50% are well tolerated and do not lead to structural or functional impairments in mouse models^19,32–35^, although other authors reported that global tau reduction might have detrimental effects in brain function ^36–39^. A recent phase I clinical trial using antisense oligonucleotides (ASOs) targeting tau, have demonstrated the feasibility and safety of *MAPT* suppression in humans^40^. However, challenges remain, including the need for repeated intrathecal administration and uncertainty about the long-term effects of sustained global tau reduction in the human brain.

In this context, endogenous RNA-targeting using artificial microRNAs (amiRNAs) offer several advantages because they can be designed using endogenous miRNA backbones to achieve high specificity for *MAPT* mRNA and their integration into cellular machinery allows for long-lasting and spatially restricted expression in target regions, potentially reducing the need for repeated dosing and minimizing off-target effects^41–44^.

While this study demonstrates the efficacy of Tau-miRNA treatment, it also presents limitations. The intervention was restricted to a single brain region (mPFC) and analyzed over a six-month period. Thus, questions remain about the durability and generalizability of the therapeutic effects. Future studies should assess long-term behavioral and circuit-level outcomes, evaluate systemic or multi-site delivery, and perform transcriptomic/proteomic analyses to understand broader molecular consequences. In addition, exploring interactions between tau reduction and other pathological pathways—such as inflammation, autophagy, or mitochondrial dysfunction—may reveal new combinatorial strategies to halt or reverse neurodegeneration.

## MATERIALS AND METHODS

### Mice and experimental design

All animal procedures were designed in accordance with the NIH Guidelines for the Care and Use of Laboratory Animals. Protocols were approved by Institutional Animal Care and Use Committee of INGEBI-CONICET. Mice were housed in standard conditions under 12 h dark/light cycle with *ad libitum* access to food and water. Htau transgenic mice ^45^ in a C57BL/6J background, were obtained from Jackson Laboratories (Bar Harbour, Maine, United States; B6.Cg-Mapt^tm1(EGFP)Klt^Tg(MAPT)8cPdav/J. Strain number: 005491) and bred in house. To confirm the presence of the human *MAPT* transgene and the mouse *Mapt*^−/−^ background, all mice used in this study were genotyped by PCR as previously described ^22^Htau mice were in-house backcrossed to C5BL/6J mice every 12 generations to refresh breeders.

Experimental groups (htau or WT littermates) at 6 months-old were randomly allocated to receive either Tau-miRNA (166+724; 1:1) or Scr-miRNA as previously described ^21^. At 12 months of age, behavioural tests and electrophysiological *in vivo* recordings were performed as previously described^21,24^. For protein extraction, mice were sacrificed by cervical dislocation and the injected area was dissected and stored at -80°C until use. For immunofluorescence array tomography analyses, mice were perfused transcardially with 4% paraformaldehyde in 0.1 M phosphate buffer (pH 7.4).

### Design of amiRNAs and subcloning in viral vectors

Artificial miRNAs were designed following described rules for siRNA^46,47^ following an in house method described^21,48^. . Target regions were chosen at Exons 2/3 junction (alternative exons) and at Exon 11 (constitutive exon) to maximize silencing. Thermodynamic values of the duplex were calculated according to the energy at the 5’ end of the sense strand (Es) and the energy of the 5’ end on the strand considered antisense (Eas). Only Eas < Es were selected with ΔT values between -5 and -3. Five initial sequences were obtained targeting the human *MAPT* transcript, of which 2 were selected after BLAST against human and mouse (sequences with >17 nt off target match were discarded). These conservative selection reduces the off target effect pf miRNAS^41,44^ The Scr sequence was obtained with the same GC content with less than 15 nt match obtained after BLAST. Selected siRNA sequences were embedded into a microRNA backbone containing the arms of miR--155 followed by the antisense sequence, the loop, and the sense sequence. From the siRNA sense sequence, nt 10 and 11 were removed to allow the formation of a 3D structure that optimizes the binding with the RISC system ^46^. Full sequences of the artificial miRNAs used in this study are as follows: [miRNA5’arm -<u>Antisense</u>(21nt) - *Loop*-<u>sense</u> (19nt)-miRNA 3’arm].

#### Tau-miRNA 166 (target E2/3)

ACCGGTGTCGACTTTAAAGGGAGGTAGTGAGTGGACCAGTGGATCCTGGAGGCTTGCTGAAGG CTGTATGCTG<u>AATGCCTGCTTCTTCAGCTTT</u>*GTTTTGGCCACTGACTGAC*<u>AAAGCTGAAAGCAGG</u>

<u>CATT</u>CAGGACACAAGGCCTGTTACTAGCACTCACATGGAACAAATGGCCCAGATCTGGCCGCACT CGAGATATCTAGAATTCACTAGTGAGCTC

#### Tau-miRNA 724 (target E11)

ACCGGTGTCGACTTTAAAGGGAGGTAGTGAGTGGACCAGTGGATCCTGGAGGCTTGCTGAAGG CTGTATGCTG<u>TAATGAGCCACACTTGGAGGT</u>*GTTTTGGCCACTGACTGAC*<u>ACCTCCAAGTGGCTC ATTA</u>CAGGACACAAGGCCTGTTACTAGCACTCACATGGAACAAATGGCCCAGATCTGGCCGCACT CGAGATATCTAGAATTCACTAGTGAGCTC

#### Scr-miRNA

accggtGTCGACTTTAAAGGGAGGTAGTGAGTGGACCAGTGGATCCTGGAGGCTTGCTGAAGGCT GTATGCTGAAATGTACTGCGCGTGGAGAC***GTTTTGGCCACTGACTGAC***<u>GTCTCCACGCAGTACA TTT</u>**CAGGA**CACAAGGCCTGTTACTAGCACTCACATGGAACAAATGGCCCAGATCTGGCCGCACT CGAGATATCTAGAATTCACTAGTGAGCTC

Artificial miRNAs were subcloned under the human synapsin promoter between AgeI and EcoRI sites into a lentiviral vector backbone previously described (REFs). Lentiviral particles were generated as previously described ^21^. Briefly, HEK-293T cells were grown on DMEM, supplemented with 10% (v/v) fetal bovine serum (FBS; Natocor, Argentina), 0.5 mM L-glutamine, 100 U/ml penicillin and 100 μg/ml streptomycin (ThermoFisher). Cells at 80-85% confluence were co-transfected with a lentiviral shuttle vector (Tau-miRNA 166, Tau-miRNA 724 or Scr-miRNA) together with helper vectors encoding packaging and envelope proteins (CMVΔ8.9 and CMV-VSVg, respectively). Viral particles were harvested from the culture medium 36h after transfection and treated with RNase-free DNase I (Thermo Fisher). Viral vectors were purified by centrifugation and filtering (45 μm pore), concentrated by ultracentrifugation at 100,000 x g (Ti 90 rotor, Beckman) and resuspended in sterile PBS. After performing titration, 10 μl aliquots of viral particles were stored at -80°C.

### Stereotaxic injections

LVs were delivered into the medial prefrontal cortex (mPFC) as previously described ^21,24^. Briefly, mice (males and females) aged 22-26 weeks (weight 25-32g) were anesthetized with isoflurane 0.5-2,5% (2,5% for induction/0.5-1% for maintenance, Baxter) in medical grade oxygen with an air flow at 2.5 L min-1 and placed into a stereotactic frame (Stoelting CO.). A 10 μL Hamilton syringe coupled to a 36G stainless steel tube (Cooper needleworks, United Kingdom) was used to inject 1.5 μL of lentiviral suspension (0.5 ×107 TU/ml; 0.2 μL/min) per site of injection, bilaterally, at 4 sites into the mPFC, following coordinates of mouse atlas (Paxinos and Franklin, 2013) (in mm): AP= +2.3, LM= ±0.5, DV= -1.8 and -2.2. One hour before surgery, mice received subcutaneous injection of systemic analgesic (Meloxicam;) which was repeated 24h and 48h after surgery. Local analgesic was injected before surgery (Lidocaine). Any animal showing signs of pain or discomfort after surgery was sacrificed following the end point protocol.

### Protein extraction and Western blotting

mPFC was dissected and homogenized with a buffer containing 50 mM Tris-HCl (pH 7.4), 150 mM NaCl, 2mM EGTA and proteases and phosphatases inhibitor cocktail (Thermo Fisher). Tissue disruption was performed with a motorized tissue grinder followed by sonication and protein extracts were centrifuged at 13,500 rpm for 15 min at 4ºC. Western blotting was performed as previously described^21^. Equal amounts of total protein (determined with Pierce BCA Protein Assay Kit, Thermo Fisher) were separated on 10-12% SDS-Polyacrylamide gels (prepared with Acrylamide and N,N′-Methylenebisacrylamide 30%) and transferred using a semi-dry transfer system to nitrocellulose membranes (BioRad). See-Blue Plus 2 (Thermo Fisher) was used as a molecular-weight marker. Membranes were blocked in 5% (w/v) non-fat dry milk (La Serenisima, Argentina), 0.05% v/v Tween 20 in TBS for 1 h at room temperature. Primary antibodies were used in diluted blocking solution to incubate blots overnight at 4°C: anti-total tau (1:10000; rabbit polyclonal; Dako, Denmark) and anti-β-actin-HRP conjugated (1:10000; mouse monoclonal; Sigma-Aldrich). After washing 3 times in TBS containing 0.05% v/v Tween 20, blots were incubated with secondary antibody goat anti-rabbit-HRP conjugated (1:2000; Thermo Fisher) for 2 h at room temperature. Proteins were visualized using ECL reagent (Thermo Fisher) exposing membranes on the GenegnomeXRQ (Syngene). Optical density was quantified using FluorChem software (Alpha Innotech) and total tau contents were normalized to actin, used as a loading control.

### Sarkosyl insolubility assay

Fractionation of insoluble proteins was performed using 250 ug of mPFC protein extract, which was incubated with 1% sarkosyl reagent (Sigma-Aldrich) for 1 h in minimal agitation at room temperature, following protocol previously reported^22^. Protein extract with 1% of sarkosyl reagent was ultracentrifuged at 39,000 rpm (1 h) at 20°C to obtain the pellet and then washed with sarkosyl 1% by ultracentrifugation at the same conditions for 15 min. Pellet was resuspended for 1 h at room temperature for Western blotting.

### In vivo electrophysiological recordings

#### Data acquisition

Electrodes for extracellular recording were made as previously described ^21^. Mice were deeply anesthetized with isoflurane (2% for induction, and 0.5-1% for maintenance, Baxter) and placed into a stereotaxic frame. The skull was exposed to clearly locate Bregma. A craniotomy was performed over the mPFC coordinates (AP= +2.1 mm, LM= ±0.5 mm, Bregma as reference). The tetrodes were lowered inside the brain at a speed rate of 10-20 μm/sec. Stable spontaneous action potentials were sought up between -1 and -2.5 mm from the surface. Electrophysiological data was recorded at different positions (up to 7 recordings were acquired per animal), with durations ranging from 15 to 30 min each.

#### Data processing and analysis

Analysis and statistical tests were implemented in MATLAB (The MathWorks Inc., USA). Raw signals were band filtered between 300 Hz and 6000 Hz. Spikes sorting was performed as follows. An automatic threshold was set at five times the standard deviation above the mean to detect the spike events. Detected spikes were partitioned into many clusters with a *k-means* method, and then were aggregated according to their interface energy for each pair. Clusters were manually split and merged according to their principal components for the subsequent analysis. Single units were classified into two groups based on their mean spike-width or waveform, measured as the time from the trough to the next peak of the mean action potential. Units with a ‘valley-to-peak’ (y-axis in the figure) greater than 440 μs were considered as putative pyramidal neurons (Figure S4A; red dots) and otherwise as putative interneurons (FigureS4A; blue dots). In this way, we distinguished neurons based on their average spike waveform, regardless of their firing rate. Rasters at 1 ms resolution containing a sequence of zeros (no spike event) and ones (spike event) were constructed for each isolated unit. On these rasters, the firing rate (spikes per second) and the interspike intervals (ISIs) were computed for each neuron. To measure the degree of burstiness in the firing of a given neuron, an autocorrelogram with time shifts ranging from 0 to +50 ms was computed from the rasters. Next, the time shift Δt at which half the accumulated autocorrelogram value was reached. Burstiness index was defined as 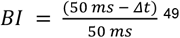.

### High-resolution immunofluorescent array tomography

After transcardiac perfusion of mice, brains were post-fixed in 4% paraformaldehyde overnight at 4°C, incubated in 15% sucrose for 24 h and then in 30% sucrose for another 24 h. Tissue blocks containing the mPFC were cut to 300 μm thick sections using a vibratome and then processed for array tomography as previously described^50^, using LRWhite resin (medium grade, Ted Pella, USA). Embedded tissue was cut with a Jumbo Histo Diamond Knife (Diatome) in an ultramicrotome (Reichert-Jung, Germany). Series of 20-30 200 nm thick sections were collected in ribbons onto glass coverslips and processed for immunofluorescence. Antibodies anti-p-tau (Thr231) AT180 (1:100; mouse; Thermo Fisher) and anti-Synapsin-1a (1:200; rabbit; Cell Signaling Technology) were used. Fluorescent-conjugated secondary antibodies raised in donkey (Alexa 488, Alexa 647, and CY3, 1:100; Jackson ImmunoResearch, United Kingdom) were used. Sections were mounted on glass slides with SlowFade Gold Antifade (Life Technologies), and then imaged in a Leica DMR fluorescence microscope using a PL APO 63X NA=1.32 oil objective and a Retiga R1 camera (Q-Imaging, United Kingdom). Serial images were aligned and converted into stacks using Fiji. A sampling mask of 120 μm X 120 μm in the mPFC was used for quantitative analysis using the Analyze Particles function, yielding values of puncta (for Synapsin) or clusters (for AT180) per μm^3^.

### Behavioral tests

Mice tested were sibling cohorts of 6 or 12 months old depending on the experimental design. Novel Object Recognition, Open Field and Elevated Plus Maze tasks were performed as described previously ^21,22,24,25^. Experiments were conducted between 13:00 h and 17:00 h under dim illumination, in a separated behavioral room, where mice were transferred in advance. Recordings were analyzed by ANY-maze (Stoelting Co.). All arenas and devices were cleaned between subjects to minimize odor cues.

### Statistical analyses

Data were analysed with Prism GraphPad software. Data sets of each experiment were classified according to the *p*-value obtained for the Shapiro-Wilk test, which determines normality. If the data set passed the test (*p*-value>0,05) the data structure was classified as Normal Distribution, and if the data set did not pass the test (*p*-value<0,05) the data structure was classified as Non-normal Distribution. For data sets classified as Normal Distribution, statistical tests used for comparing groups depended on the number of groups and independent variables used in the experiment: Unpaired t-test (two groups, one independent variable), One-way ANOVA test (three or more groups, one independent variable) or Two-way ANOVA test (two independent variables). For One-way ANOVA, *post hoc* tests were used according to the type of comparisons between groups that were relevant for the experiment: Tukey’s *post hoc* test (for comparing the mean of each group with the mean of every other group) and Dunnett’s *post hoc* test (for comparing the mean of each group with the mean of a control group). When data sets had Non-normal Distribution, statistical tests used for comparing groups were the non-parametric tests: Mann-Whitney U test (comparisons between two groups, one independent variable) or Two-sample Kolmogorov-Smirnov test (for comparisons of cumulative distribution of the data sets).

## FIGURE LEGENDS

**Figure S1. Tau local reduction by Tau-miRNA does not alter anxiety and exploratory behaviours, related to Figure 1. A-B**. Total exploration time in the Novel Object Recognition (NOR) task for each miRNA-injected group, (**A**) analysed at 12 months of age and (**B**) comparing each animal before (Pre-inj 6m) and after (Post-inj 12m) the injection. For Figure A: WT Scr-miRNA=18, WT Tau-miRNA=13, htau Scr-miRNA=14 and htau Tau-miRNA=17; and B: WT Scr-miRNA=8, WT Tau-miRNA=4, htau Scr-miRNA=7, htau Tau-miRNA=9. **C**. Discrimination index in the NOR task separated between females (F) and males (M). WT Scr-miRNA: (F=11 and M=7), WT Tau-miRNA: (F=9 and M=4), htau Scr-miRNA (F=9 and M=5) and htau Tau-miRNA (F=8 and M=9). **D-F**. Elevated Plus Maze test. **D**. Representative traces of exploration for each injected group, where sections filled in pink are the open arms and the empty ones are the closed arms. Time spent in the open arms relative to total time of exploration for each group, (**E**) assessed at 12 months and (**F**) before and after treatment. For Figure E: WT Scr-miRNA=18, WT Tau-miRNA=10, htau Scr-miRNA=12 and htau Tau-miRNA=15; and F: WT Scr-miRNA=5, WT Tau-miRNA=5, htau Scr-miRNA=6, htau Tau-miRNA=5. For Figures A, C and E data is shown as scatter dot plots with bars ± SEM and statistical analyses were performed by One-way ANOVA followed by Tukey’s *post hoc* test (A and E) and Student’s *t*-test (C). For figures B and F, data is shown as scatter dot plots, paired *t*-test.

**Figure S2. Tau knockdown by Tau-miRNAs modulates the patterning activity of cortical neurons in the PFC of htau mice, related to Figure 2. A**. Raster plot showing valley to peak and half amplitude duration of the mean spike waveform from each recorded neuron with a signal-to-noise ratio above 5. These features of spike waveforms sorted neurons in two clusters, corresponding to putative pyramidal neurons (pPYR, red dots) and putative interneurons (pINT, blue dots). **B-D**. ISI histograms plotted on a log scale for a single representative neuron from WT Scr-(**B**), htau Scr-(**C**) and htau Tau-(**D**) miRNA injected groups.

**Figure S3. Reduction of total, p-tau and insoluble tau protein contents in the PFC of injected mice, related to Figure 3**. Full bots used for quantification of (**A**) total tau protein, (**B**) p-tau and (**C**) insoluble tau contents in the mPFC of miRNAs-injected mice. For total tau and p-tau quantification, actin was used as a loading control and tau KO as a negative control. (*) indicate outlier samples not included in the analyses. Red arrows show the samples used for the panels shown in the main section.

## REFERENCES

1. Spillantini, M. G. & Goedert, M. Tau pathology and neurodegeneration. Lancet Neurol. 12, 609–622 (2013).

2. Guo, T., Noble, W. & Hanger, D. P. Roles of tau protein in health and disease. Acta Neuropathologica vol. 133 665–704 (2017).

3. Morris, M., Maeda, S., Vossel, K. & Mucke, L. The many faces of tau. Neuron 70, 410–426 (2011).

4. Brandt, R. & Götz, J. Special issue on ‘Cytoskeletal proteins in health and neurodegenerative disease’. Brain Res. Bull. 126, 213–216 (2016).

5. Jadhav, S. et al. A walk through tau therapeutic strategies. Acta Neuropathol. Commun. 7, 22 (2019).

6. Dujardin, S. et al. Different tau species lead to heterogeneous tau pathology propagation and misfolding. Acta Neuropathol. Commun. 6, 132 (2018).

7. Colom-Cadena, M. et al. Synaptic oligomeric tau in Alzheimer’s disease - A potential culprit in the spread of tau pathology through the brain. Neuron (2023) doi:10.1016/J.NEURON.2023.04.020.

8. Zempel, H. & Mandelkow, E. Lost after translation: Missorting of Tau protein and consequences for Alzheimer disease. Trends Neurosci. 37, 721–732 (2014).

9. Di Xia, Gutmann, J. M. & Götz, J. Mobility and subcellular localization of endogenous, gene-edited Tau differs from that of over-expressed human wild-type and P301L mutant Tau. Sci. Rep. 6, 29074 (2016).

10. Buée, L. et al. From tau phosphorylation to tau aggregation: what about neuronal death? Biochem. Soc. Trans. 38, 967–972 (2010).

11. Crimins, J. L., Rocher, A. B. & Luebke, J. I. Electrophysiological changes precede morphological changes to frontal cortical pyramidal neurons in the rTg4510 mouse model of progressive tauopathy. Acta Neuropathol. 124, 777–95 (2012).

12. Braak, H. & Braak, E. Neuropathological stageing of Alzheimer-related changes. Acta Neuropathol. 82, 239–259 (1991).

13. Lasagna-Reeves, C. A. et al. Tau oligomers impair memory and induce synaptic and mitochondrial dysfunction in wild-type mice. Mol. Neurodegener. 6, 39 (2011).

14. Congdon, E. E. & Duff, K. E. Is tau aggregation toxic or protective? Journal of Alzheimer’s Disease vol. 14 453–457 (2008).

15. Lane-Donovan, C. & Boxer, A. L. Disentangling tau: One protein, many therapeutic approaches. Neurotherapeutics 21, e00321 (2024).

16. Rösler, T. W., Costa, M. & Höglinger, G. U. Disease-modifying strategies in primary tauopathies. Neuropharmacology (2019) doi:10.1016/j.neuropharm.2019.107842.

17. Wegmann, S. et al. Persistent repression of tau in the brain using engineered zinc finger protein transcription factors. Sci. Adv. 7, eabe1611 (2021).

18. Shin, M.-K. et al. Reducing acetylated tau is neuroprotective in brain injury. Cell 0, (2021).

19. DeVos, S. L. et al. Tau reduction prevents neuronal loss and reverses pathological tau deposition and seeding in mice with tauopathy. Sci. Transl. Med. 9, (2017).

20. Albert, M. et al. Prevention of tau seeding and propagation by immunotherapy with a central tau epitope antibody. Brain 142, 1736–1750 (2019).

21. Facal, C. L. et al. Tau reduction with artificial microRNAs modulates neuronal physiology and improves tauopathy phenotypes in mice. Mol. Ther. 32, 1080–1095 (2024).

22. Espíndola, S. L. et al. Modulation of Tau Isoforms Imbalance Precludes Tau Pathology and Cognitive Decline in a Mouse Model of Tauopathy. Cell Rep. 23, 709–715 (2018).

23. Polydoro, M., Acker, C. M., Duff, K., Castillo, P. E. & Davies, P. Age-dependent impairment of cognitive and synaptic function in the htau mouse model of tau pathology. J. Neurosci. 29, 10741–10749 (2009).

24. Muñiz, J. A. et al. SMaRT modulation of tau isoforms rescues cognitive and motor impairments in a preclinical model of tauopathy. Front. Bioeng. Biotechnol. 10, 1–13 (2022).

25. Damianich, A. et al. Tau mis-splicing correlates with motor impairments and striatal dysfunction in a model of tauopathy. Brain 144, 2302–2309 (2021).

26. Ittner, L. M. et al. Dendritic function of tau mediates amyloid-β toxicity in alzheimer’s disease mouse models. Cell 142, 387–397 (2010).

27. Bi, M. et al. Tau exacerbates excitotoxic brain damage in an animal model of stroke. Nat. Commun. 8, (2017).

28. Roberson, E. D. et al. Reducing endogenous tau ameliorates amyloid beta-induced deficits in an Alzheimer’s disease mouse model. Science 316, 750–754 (2007).

29. Wu, J. W. et al. Neuronal activity enhances tau propagation and tau pathology in vivo. Nat. Neurosci. 19, 1085–1092 (2016).

30. Chang, C. W., Shao, E. & Mucke, L. Tau: enabler of diverse brain disorders and target of rapidly evolving therapeutic strategies. Science 371, (2021).

31. Chang, C. W., Evans, M. D., Yu, X., Yu, G. Q. & Mucke, L. Tau reduction affects excitatory and inhibitory neurons differently, reduces excitation/inhibition ratios, and counteracts network hypersynchrony. Cell Rep. 37, (2021).

32. Shao, A. E., Chang, C., Li, Z., Yu, X. & Ho, K. Tau ablation in excitatory neurons and postnatal tau knockdown reduce epilepsy, SUDEP, and autism behaviors in a Dravet syndrome model. Sci. Transl. Med. 5527, (2022).

33. Chang, C. W., Evans, M. D., Yu, X., Yu, G. Q. & Mucke, L. Tau reduction affects excitatory and inhibitory neurons differently, reduces excitation/inhibition ratios, and counteracts network hypersynchrony. Cell Rep. 37, (2021).

34. Götz, J.Bodea, L.-G. & Goedert, M. Rodent models for Alzheimer disease. Nat. Rev. Neurosci. (2018) doi:10.1038/s41583-018-0054-8.

35. Wegmann, S. et al. Removing endogenous tau does not prevent tau propagation yet reduces its neurotoxicity. EMBO J. 1–14 (2015) doi:10.15252/embj.201592748.

36. Ma, Q.-L. et al. Loss of MAP function leads to hippocampal synapse loss and deficits in the Morris Water Maze with aging. J. Neurosci. 34, 7124–36 (2014).

37. Chang, C.-W. et al. Tau reduction affects excitatory and inhibitory neurons differently, reduces excitation/inhibition ratios, and counteracts network hypersynchrony HHS Public Access In brief. Cell Rep 37, 109855 (2021).

38. Avila, J. Our Working Point of View of Tau Protein. Journal of Alzheimer’s Disease vol. 62 1277–1285 (2018).

39. Maya-Monteiro, C. M. et al. Behavioral Abnormalities in Knockout and Humanized Tau Mice. Front. Endocrinol. | https://www.frontiersin.org 1, p124 (2020).

40. Mummery, C. J. et al. Tau-targeting antisense oligonucleotide MAPTRx in mild Alzheimer’s disease: a phase 1b, randomized, placebo-controlled trial. Nat. Med. (2023) doi:10.1038/s41591-023-02326-3.

41. Fowler, D. K., Williams, C., Gerritsen, A. T. & Washbourne, P. Improved knockdown from artificial microRNAs in an enhanced miR-155 backbone: A designer’s guide to potent multi-target RNAi. Nucleic Acids Res. 44, e48 (2015).

42. Chen, S. K. et al. Efficacy and safety of a SOD1-targeting artificial miRNA delivered by AAV9 in mice are impacted by miRNA scaffold selection. Mol. Ther. Nucleic Acids 34, (2023).

43. Diener, C., Keller, A. & Meese, E. Emerging concepts of miRNA therapeutics: from cells to clinic. Trends Genet. 38, 613–626 (2022).

44. Kotowska-Zimmer, A., Pewinska, M. & Olejniczak, M. Artificial miRNAs as therapeutic tools: Challenges and opportunities. Wiley Interdiscip. Rev. RNA 12, e1640 (2021).

45. Andorfer, C. et al. Hyperphosphorylation and aggregation of tau in mice expressing normal human tau isoforms. J Neurochem 86, 582–590 (2003).

46. Pei, Y. & Tuschl, T. On the art of identifying effective and specific siRNAs. Nat. Methods 3, 670–676 (2006).

47. Tafer, H. Bioinformatics of siRNA design. Methods Mol. Biol. 1097, 477–490 (2014).

48. Bordone, M. P. et al. Fyn knockdown prevents levodopa-induced dyskinesia in a mouse model of Parkinson’s disease. eNeuro 8, (2021).

49. Csicsvari, J., Hirase, H., Czurko, A. & Buzsáki, G. Reliability and State Dependence of Pyramidal Cell–Interneuron Synapses in the Hippocampus. Neuron 21, 179–189 (1998).

50. Soiza-Reilly, M. et al. SSRIs target prefrontal to raphe circuits during development modulating synaptic connectivity and emotional behavior. Mol. Psychiatry 24, 726–745 (2019).

