## Supplementary figures and images for "Artificial microRNAs targeting Tau enable post-symptomatic functional recovery in aged tauopathy mice"

### Supplemental Figure 1

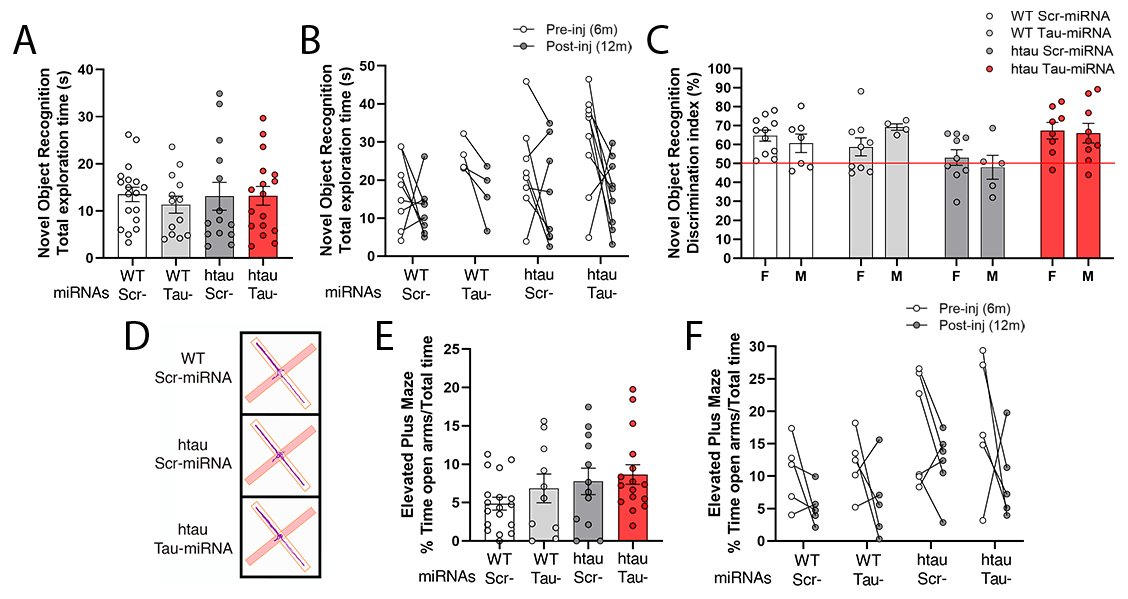

### Supplemental Figure 2

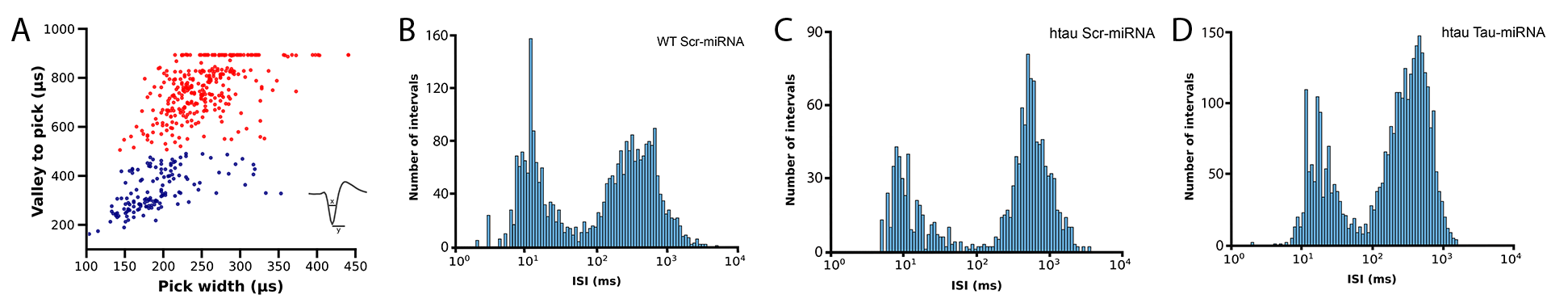

### Supplemental Figure 3

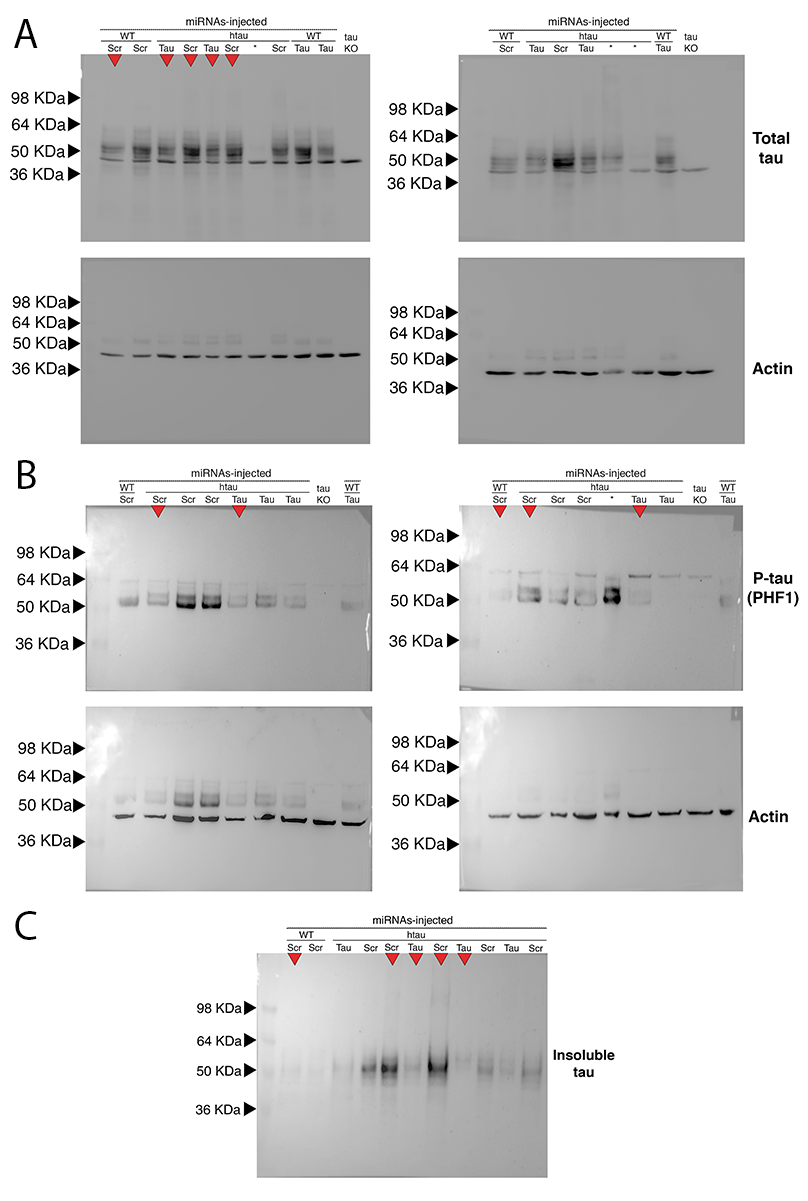
